# Epstein-Barr virus induced 3 attributes to TLR7-mediated splenomegaly and bicytopenia

**DOI:** 10.1101/2024.06.10.598101

**Authors:** Masanori Iseki, Yuma Sakamoto, Daiki Takezaki, Yoshihiro Matsuda, Mariko Inoue, Shin Morizane, Tomoyuki Mukai

**Author notes:** Correspondence: Tomoyuki Mukai, M.D., Ph.D., Department of Immunology and Molecular Genetics, Kawasaki Medical School 577 Matsushima, Kurashiki, Okayama 701-0192, Japan.

## Abstract

Epstein-Barr virus induced 3 (EBI3) is a gene induced by stimulation of toll-like receptors (TLRs) and functions as a component of a heterodimer cytokine IL-27. IL-27 regulates both innate and acquired immune responses; however, their function *in vivo* is still largely unknown. Splenomegaly, an enlargement of the spleen, is known to be induced by chronic infectious diseases, including infectious mononucleosis due to EB virus infection. Repeated treatment of imiquimod (IMQ; a TLR7 agonist) has been reported to induce splenomegaly and cytopenia due to increased splenic function. Although immune cell activation is speculated to be involved in the pathogenesis of chronic infection-mediated splenomegaly, the detailed molecular mechanism is unknown. Here, we demonstrated that IMQ induced marked splenomegaly and severe bicytopenia (anemia and thrombocytopenia) in the wild-type mice. Myeloid cells, not lymphoid cells, were increased in the enlarged spleen. Extramedullary hematopoiesis was observed in the enlarged spleen of the IMQ-treated mice. RNA-seq analysis revealed that type I interferon (IFN)-related genes were upregulated in the spleen and peripheral blood in IMQ-treated mice. *Ebi3* deficiency partially retrieved these IMQ-induced pathological changes. We found that *Il27* as well as *Ebi3* genes were elevated by IMQ in the spleen and peripheral blood. Further, IL-27 stimulation upregulated type I IFN-related genes in bone marrow-derived macrophage culture in the absence of type I IFN. Collectively, EBI3 contributes to TLR7-induced splenomegaly and bicytopenia, presumably via IL-27.

## Introduction

Epstein-Barr virus induced 3 (EBI3) is a gene that is originally reported to be upregulated in B cell lines infected with EB virus. EBI3 is expressed in a variety of immune cells (1–3), and its expression is induced by stimulation of Toll-like receptors (TLRs) (4). It binds to IL-27p28 and functions as a heterodimer cytokine named IL-27 (5). IL-27 is a cytokine that functions in both innate and acquired immunity. Human monocytes stimulated with IL-27 upregulate the expression of pro-inflammatory cytokine genes such as *IL1B*, *TNF*, *IL12A*, and *IL18* (6, 7). IL-27 has been reported to act directly on CD4^+^ T cells to promote differentiation into Th1 cells while inhibiting differentiation into Th2 and Th17 cells (8–10). This cytokine directly affects hematopoietic stem cells (HSCs) and promotes their differentiation into myeloid cells (11, 12). Additionally, recent studies have indicated that IL-27 induces the expression of type I IFN-related genes independently of type I IFN, which plays a role in defense mechanisms against viral infection (13).

Spleen is an essential organ for immune defense against blood-borne antigens (14, 15). Splenomegaly, an enlargement of the spleen, is caused by chronic infectious diseases and leukemia, leading to a serious anemia and thrombocytopenia due to splenic hyperfunction (16, 17). Additionally, splenomegaly due to infectious mononucleosis (EB virus infection) has been reported to be fatal due to splenic rupture (18).

TLR7 recognizes single-stranded RNAs from bacteria and viruses (19–21). Plasmacytoid dendritic cells (pDCs), which highly express the TLR7, produce large amounts of type-I interferons (IFNs, IFN-α/β), which in turn activate various immune cells and act as an immune defense (19–26). Sustained TLR7 stimulation has been applied as a murine model to investigate the innate and acquired immune systems (27–30). Repeated treatment of imiquimod (IMQ; a TLR7 agonist) has been reported to induce splenomegaly and cytopenia due to increased splenic function (27, 29). Although immune cell activation is speculated to be involved in the pathogenesis of chronic infection-mediated splenomegaly, the detailed molecular mechanism is unknown.

In this study, we investigated the role of EBI3 in TLR7-induced splenomegaly and cytopenia. To this end, we applied IMQ to the wild-type (WT) mice and *Ebi3* gene-deficient (KO) mice. We found that IMQ induced severe splenomegaly and bicytopenia in the WT mice, which was associated with increased myeloid cells in the spleen and increased type I IFN-related genes in the spleen. *Ebi3* deficiency reduced the IMQ-mediated pathological changes. IL27, a heterodimer of EBI3 and IL-27p28, induced type I IFN-related genes, independently of type I IFN. Based on the findings, we presume that EBI3 is involved in the IMQ-induced splenomegaly and bicytopenia, presumably via IL-27.

## Methods

### Antibodies and reagents

Anti-EBI3 (clone EPR28427-89, Abcam, Cambridge, UK), anti-IL27A (clone EPR18247-86, Abcam), anti-actin antibody (A2066, Sigma-Aldrich, St. Louis, MO, USA), and horseradish peroxidase-conjugated anti-rabbit IgG antibody (7074, Cell Signaling Technology, Danvers, MA, USA) were used for Western blotting. Anti-Factor VIII (clone EPR24039-262, Abcam) antibody was used for Immunohistochemistry. The following antibodies were used for flow cytometry: Brilliant Violet 510-conjugated anti-CD19 (clone RA3-6B2, BioLegend, San Diego, CA, USA), phycoerythrin (PE)-Cy7-conjugated anti-CD3ε (clone 145-2C11, BioLegend), violetFluor 450-conjugated anti-CD4 (clone RM4-5, Tonbo Biosciences, San Diego, CA, USA), allophycocyanin (APC)-conjugated anti-CD8α (clone 53-6.7, BioLegend), FITC-conjugated anti-CD44 (clone IM7, BioLegend), PE-conjugated anti-CD62L (clone MEL-14, Tonbo Biosciences), APC-conjugated anti-CD11c (clone N418, BioLegend), Pacific Blue-conjugated anti-B220/CD45R (clone RA3-6B2, BioLegend), APC-Cy7-conjugated anti-MHC class II (I-A/I-E) (clone M5/114.15.2, BioLegend), Pacific Blue-conjugated anti-CD11b (clone M1/70, BioLegend), FITC-conjugated anti-Gr-1 (clone RB6-8C5, BioLegend), and APC-conjugated anti-TER-119 (clone TER-119, BioLegend). To detect dead cells, 7-amino-actinomycin D (7AAD, AAT Bioquest, Pleasanton, CA, USA) was used.

### Generation of *Ebi3*-deficient mice

*Ebi3*-deficient (KO) mice were generated at Kawasaki Medical School (Kurashiki, Japan). The mice were generated by clustered regularly interspaced short palindromic repeats (CRISPR)/Cas9-mediated deletion of the exon 2 of the EBI3 coding sequence using genome editing by electroporation of Cas9 protein (GEEP) methods (31, 32). CRISPR RNAs (crRNAs) were designed as follows: gRNA#1: 5′-TATACTTACTACAAACCCCA-3′ and gRNA#2: 5′-TCAGGGGGTGATATGCTCAG-3′ (Supplementary Fig. 1A). Synthetic trans-activating CRISPR RNA (tracrRNA) and a recombinant Cas9 protein were obtained from Integrated DNA Technologies Inc. (Coralville, IA, USA). Deletion of the target region was confirmed by PCR analysis and DNA sequencing. Off-target candidate sequences were checked using COSMID (CRISPR search with mismatches, insertions, and/or deletions, https://crispr.bme.gatech.edu/). We confirmed no off-target mutations in the generated KO mice among the top three candidate sequences for each guide RNA. Genotypes of the KO mice were confirmed by PCR analysis using the following primers, forward 5′-CAGTTTTCTCATCTAGAAATTGAGCTA-3′, reverse 5′-CCATGGTGAATCTCGAAAGG-3′ (Supplementary Fig. S1A and B). The lack of *Ebi3* mRNA and EBI3 protein expression was confirmed by RT-PCR and Western blotting, respectively (Supplementary Fig. 1B and C).

### Mice

WT C57BL/6J mice were purchased from CLEA Japan (Tokyo, Japan). WT and *Ebi3*-deficient mice were housed at the Kawasaki Medical School animal facility in groups (2–5 mice per cage) and maintained at 22 °C in 12 h light/12 h dark cycles with free access to water and food (MF diet, Oriental Yeast Co., Tokyo, Japan). All animal experiments were approved by the Institutional Safety Committee for Recombinant DNA Experiments (20-49, 0004-00) and the Institutional Animal Care and Use Committee of Kawasaki Medical School (22-072, 23-092). All experimental procedures were conducted according to institutional and NIH guidelines for the humane use of animals.

### IMQ treatment

*Ebi3* KO mice and sex-matched WT mice (9–12 weeks old) were treated epicutaneously with 25 mg of 5% IMQ cream (Beselna Cream, Mochida Pharmaceutical, Tokyo, Japan) on their right ear three times a week for up to 8 weeks. Vaseline was similarly applied to the ears of WT and KO mice as controls. Blood, spleen, skin, tibia, and femur samples were collected from the mice after euthanasia.

### Complete blood counts

Ethylenediaminetetraacetic acid (EDTA) was added to the fresh blood samples to a final concentration of 1 mg/ml, and the samples were mixed thoroughly. The white blood cell (WBC), red blood cell (RBC), and platelet (PLT) were subsequently counted by the automated hematology analyzer (Dri-chem 7000V-Z, Fujifilm, Tokyo, Japan).

### Flow cytometry

Flow cytometry of the spleen was performed as reported previously (33). Splenocytes were prepared by mashing the spleen with frosted glass slides. After the cells were washed, erythrocyte lysis was performed by ACK lysis buffer (155 mM NH_4_Cl, 10 mM KHCO_3_, 0.1 mM Na_2_EDTA). After Fc-blocking with rat anti-mouse CD16/CD32 (Mouse BD Fc Block, 553142, BD Biosciences), the cells were incubated in staining buffer (PBS containing 2% fetal calf serum (FCS) and 0.09% NaN_3_) with an antibody cocktail for 20 min on ice in the dark. After washing twice with staining buffer, the cells were suspended in a staining buffer containing 0.5 μg/ml of 7AAD. Data were acquired using a FACSCanto II (BD Biosciences) and analyzed with FlowJo software (Version 10, BD Biosciences). Gating strategies for the spleen cells are shown in Supplementary Figure 2.

### Histological analysis

The spleens were fixed in 4% paraformaldehyde solution for 48 hours and embedded in paraffin. The femurs were fixed in 4% paraformaldehyde solution for 48 hours, demineralized in a neutral demineralizing solution (10% EDTA, pH 7.2) for a minimum of two weeks, and then embedded in paraffin. The tissues were cut into 3-μm thin sections, and the sections were stained with hematoxylin and eosin. Digital images of the stained sections were obtained using an IX83 inverted microscope (Olympus, Tokyo, Japan).

### Bone marrow-derived pDC culture

Culture of bone marrow-derived pDC was performed according to a guideline for DC generation (34). Bone marrow cells were harvested from the femurs and tibias from WT and KO mice. Red blood cell-lysed bone marrow cells were cultured in culture medium (RPMI1640 containing 10% FCS, penicillin-streptomycin, 50 μM 2-mercaptoethanol) with 100 ng/ml of human FMS-like tyrosine kinase 3 ligand (FLT3L, PeproTech, Cranbury, NJ, USA) for 7 days. The percentage of pDCs (CD11c^+^ B220^+^) was measured by flow cytometry. The FLT3L-treated cells were plated onto 12-well culture plates at a density of 1 × 10^6^ cells/ml, allowed to rest for at least five hours, and then stimulated with IMQ (InvivoGen, San Diego, CA, USA) or CpG ODN 1826 (InvivoGen) for two or six hours. The culture supernatants were collected and stored at −80°C. RNA samples were collected using RNAiso Plus (9108, Takara Bio).

### Bone marrow-derived macrophage culture

Mouse bone marrow cells were isolated from femurs and tibias. Non-adherent bone marrow cells were seeded at a density of 1.0 × 10^6^ cells/ml on 10-cm culture dishes. The cells were incubated for 7 days in α-minimum essential medium (MEM) supplemented with 10% FCS and recombinant mouse macrophage colony stimulating factor (25 ng/ml, PeproTech) at 37°C and 5% CO_2_. The percentage of macrophages (CD11b^+^ F4/80^+^) was measured by flow cytometry, and we confirmed that the percentages were over 95% of live cells in our culture system. The macrophages were plated onto 12-well culture plates at a density of 1.6 × 10^5^ cells/ml, allowed to rest overnight, and then stimulated with recombinant mouse IL-27 (100 ng/ml, R&D Systems, Minneapolis, MN, USA) for 6 hours. RNA samples were collected.

### Real-time quantitative polymerase chain reaction

Quantitative polymerase chain reaction (qPCR) was performed as previously described (35, 36). Total RNA from cultured cells or spleens was extracted using RNAiso Plus (Takara Bio). Total RNA from mouse whole blood was isolated using PAXgene Blood RNA Tubes and kit (761265, BD Biosciences, San Jose, CA, USA; 762174, Qiagen, Hilden, Germany), as reported previously (32, 37). RNA was then solubilized in ribonuclease (RNase)-free water. Complementary DNA was synthesized using the PrimeScript RT Reagent Kit (RR037, Takara Bio). qPCR was performed using TB Green PCR Master Mix (RR820, Takara Bio) with the StepOnePlus Real-Time PCR System (Thermo Fisher Scientific, Cleveland, OH, USA). Gene expression levels were calculated relative to *Hprt* expression using the ΔΔCt method and normalized to the indicated control samples. The primers used in this study are listed in Supplementary Table 1. A mixture of four primers was used to detect all *Ifna* isoforms, as reported (38). All qPCR reactions yielded products with single-peak dissociation curves.

### Enzyme-linked immunosorbent assay

Mouse sera were collected using Bloodsepar (31204, Immuno-Biological Laboratories, Fujioka, Japan) and stored at −80°C. Concentrations of IFN-α in culture supernatants or mouse serum were measured using Mouse IFN-alpha All Subtype Quantikine ELISA Kit (MFNAS0, R&D Systems) according to the manufacturer’s protocols.

### Multiplex cytokine assay

Multiplex cytokine assay was performed as reported previously (39). Serum concentrations of 10 cytokines (IFN-γ, IL-10, IL-12p70, IL-17A, IL-1β, IL-2, IL-4, IL-6, MCP-1/CCL2, TNF-α) were quantified using the MILLIPLEX Mouse Cytokine/Chemokine Magnetic Bead Panel (Merck Millipore, Burlington, MA, USA) and Luminex 200 Instrument System (Merck Millipore), in accordance with the manufacturer’s protocol.

### Western blotting

Western blotting was performed as described (40, 41). Spleen tissues were lysed in RIPA lysis buffer (R0278; Sigma-Aldrich) containing a protease inhibitor cocktail (P8340, Sigma-Aldrich). Protein concentrations were determined using a BCA Protein Assay Kit (23227, Thermo Fisher Scientific). All samples were resolved using sodium dodecyl sulfate-polyacrylamide gel electrophoresis and transferred onto PVDF membranes. For blocking, 5% skim milk in Tris-buffered saline with 0.1% Tween-20 (TBS-T) was used. After blocking, the membranes were incubated with the indicated primary antibodies, followed by incubation with appropriate horseradish peroxidase-conjugated species-specific secondary antibodies. Bands were detected using the Clarity Max Western ECL Substrate (Bio-Rad, Hercules, CA, USA) and visualized using ImageQuant LAS-4000 (GE Healthcare, Little Chalfont, UK). Actin was used as the loading control to normalize the amount of protein.

### RNA-seq analysis

Total RNA from spleens was extracted with RNAiso Plus and cleaned up with RNeasy Mini Kit (Qiagen). Following the RNA concentration measurement by NanoDrop spectrophotometer (Thermo Fisher Scientific), pooled RNA samples were prepared by combining equal RNA amounts of each sample for each WT-Vaseline, KO-Vaseline, WT-IMQ, and KO-IMQ groups. Total RNA from mouse peripheral blood was isolated using the PAXgene Blood RNA kit. Pooled samples of blood RNA were prepared in an identical manner. RNA-seq libraries were prepared using the NEBNext Ultra II RNA Library Prep Kit for Illumina (New England Biolabs, Ipswich, MA, USA) according to the manufacturer’s protocol. Sequencing was performed using NovaSeq 6000 sequencer (Illumina, San Diego, CA). Gene ontology (GO) analysis was performed on the up- and down-regulated genes using gprofiler2 and clusterProfiler. In gprofiler2, Fisher’s exact probability test was conducted to determine which terms were biased toward differentially expressed genes. Corrected p-values were obtained by the BH method. Next, using clusterProfiler, Fisher’s exact probability test was performed. A balloon plot was created by sorting the terms in order of increasing gene ratio (count data ÷ the number of differentially expressed genes).

### Immunohistochemistry

Factor VIII expression was determined in spleen tissues by immunohistochemical staining as described previously (41). Tissue sections were deparaffinized and rehydrated. Epitope retrieval was performed by heating the slides three times for 5 min at 100 °C in 0.01 M citrate buffer (pH 6). Endogenous peroxidase activity was inhibited using 3.0% hydrogen peroxide in methanol for 10 min. After blocking with normal goat serum, the slides were incubated with an anti-Factor VIII antibody (1:1000 dilution) overnight at 4 °C, followed by 60 min of incubation with a horseradish peroxidase-conjugated goat anti-rabbit secondary antibody. Finally, the indicated protein expression was visualized using a 3,3-diaminobenzidine substrate–chromogen system (Nichirei Biosciences Inc., Tokyo, Japan). Tissue sections stained with isotype controls were used as the negative controls.

### Statistical analysis

All values are presented as mean + standard deviation. Data were analyzed on Prism version 8 (GraphPad Software, Boston, MA, USA). The Mann-Whitney test was used to compare two groups, and the Kruskal-Wallis test followed by the Dunn’s multiple comparisons test was used to compare three or more groups. The threshold for statistical significance was set at *p* < 0.05.

## Results

### Deletion of EBI3 reduces IMQ-induced splenomegaly and bicytopenia

Repeated TLR7 stimulation has been reported to induce splenomegaly and cytopenia (27–30). To clarify the involvement of EBI3 in splenomegaly and blood cell depletion under TLR7 stimulation, we applied IMQ cream to the ears of WT or *Ebi3* KO mice three times a week for 8 weeks. After 8 weeks of IMQ treatment, WT mice showed marked splenomegaly (Fig. 1A, C and D). Although there were no statistically significant differences in body weight change among the groups (Fig. 1B), splenomegaly was significantly reduced in the IMQ-applied KO mice compared to the IMQ-treated WT mice (Fig. 1A, C and D). The number of leukocytes in the peripheral blood was increased in the IMQ-treated WT mice, while the increase was not observed in IMQ-treated KO mice (Fig. 1E). Peripheral blood analysis also revealed that IMQ-applied WT exhibited severe anemia and thrombocytopenia and that these findings were partially retrieved in the KO mice (Fig. 1F-H).

**Figure 1.**
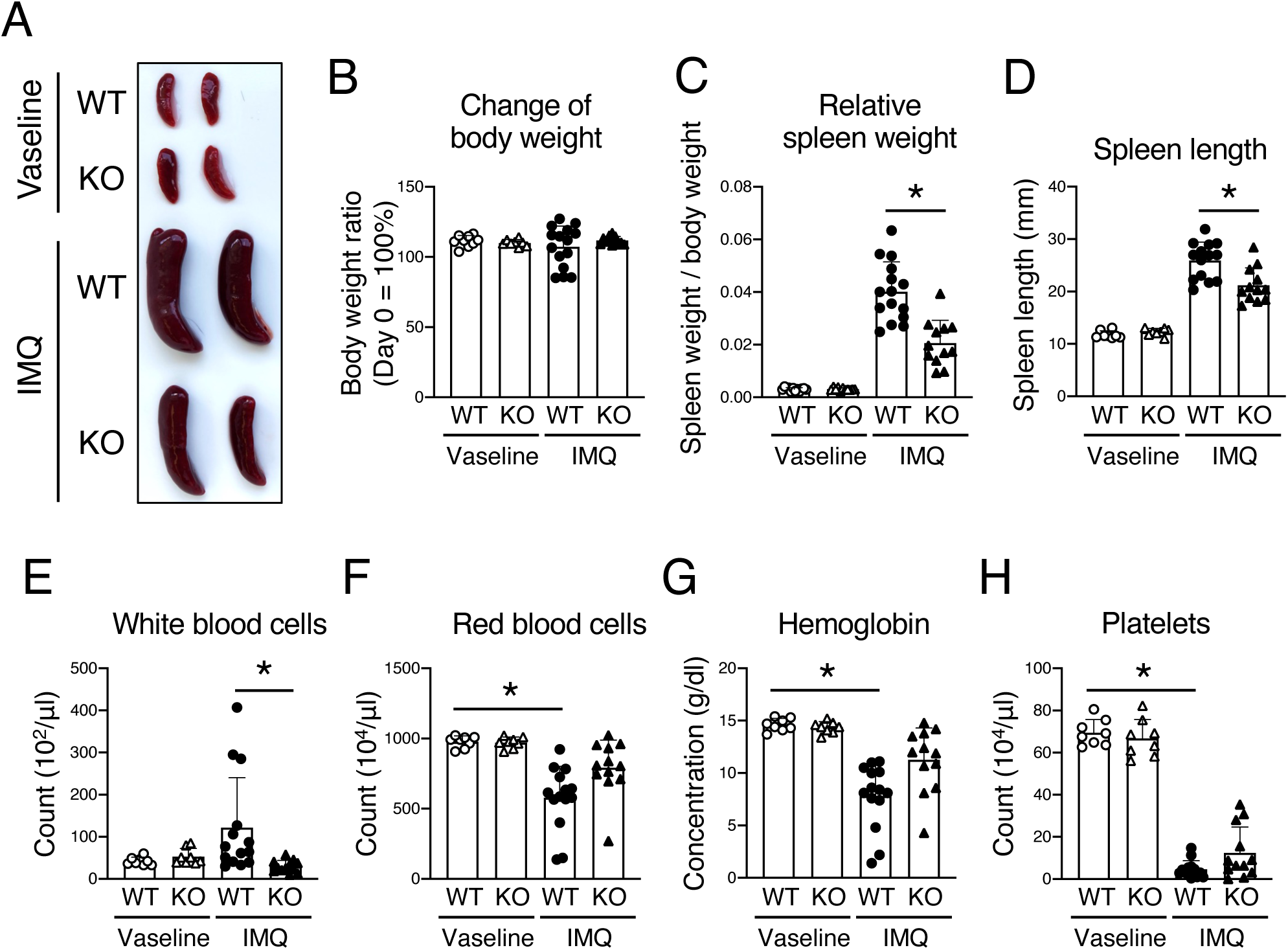
*Ebi3* deficiency retrieved imiquimod (IMQ)-induced splenomegaly and aberrant blood cell count. IMQ or Vaseline (control) was applied to the ears of WT or *Ebi3* knock-out (KO) mice three times a week for 8 weeks. (A) Representative macroscopic images of the spleens. (B) Change of body weight, relative to 100% at day 0. (C) The ratio of spleen weight relative to body weight. (D) The length of the spleens. (E-H) White blood cell count (E), red blood cell count (F), hemoglobin concentration (G), and platelet count (H) in the peripheral blood of WT or *Ebi3* KO mice. All data are expressed as the mean + standard deviation. \**P* < 0.05 by the Mann-Whitney test (C and D) or the Kruskal-Wallis test with the Dunn’s multiple comparisons test (E-H).

Analysis of mice after 2 or 4 weeks IMQ application revealed that splenomegaly progressed in proportion to the duration of IMQ application and that thrombocytopenia was observed after 4 weeks of IMQ treatment (Supplementary Fig. 3A-C). These results suggest that splenomegaly is apparent prior to anemia and thrombocytopenia. In addition, EBI3 deficiency tended to reduce the pathological changes at least 2 weeks later (Supplementary Fig. 3A-C).

### IMQ application altered immune cell subsets in the spleen, which were alleviated by EBI3 deficiency

To examine the changes in immune cell subsets in the enlarged spleen, flow cytometry analysis was performed. We found a significant decrease in the percentages of B cells and T cells and an increase in the percentages of non-T, non-B cells in the IMQ-treated WT mice compared to those in the Vaseline-treated WT mice. Although similar changes were observed in the IMQ-treated KO mice, the changes were to a lesser extent than those in IMQ-treated WT mice (Fig. 2A upper and B). Furthermore, flow cytometric analysis of non-T, non-B cells revealed that all populations, including monocytes/granulocytes (CD11b^high^ CD11c^low^), macrophages (CD11b^int^ CD11c^int^), dendritic cells (DCs, CD11b^low^ CD11c^high^), non-myeloid cells (CD11b^-^ CD11c^-^), exhibited an increased percentage in the spleen in IMQ-applied WT mice, while induction was milder in KO mice than in WT mice (Fig. 2A lower and C).

**Figure 2.**
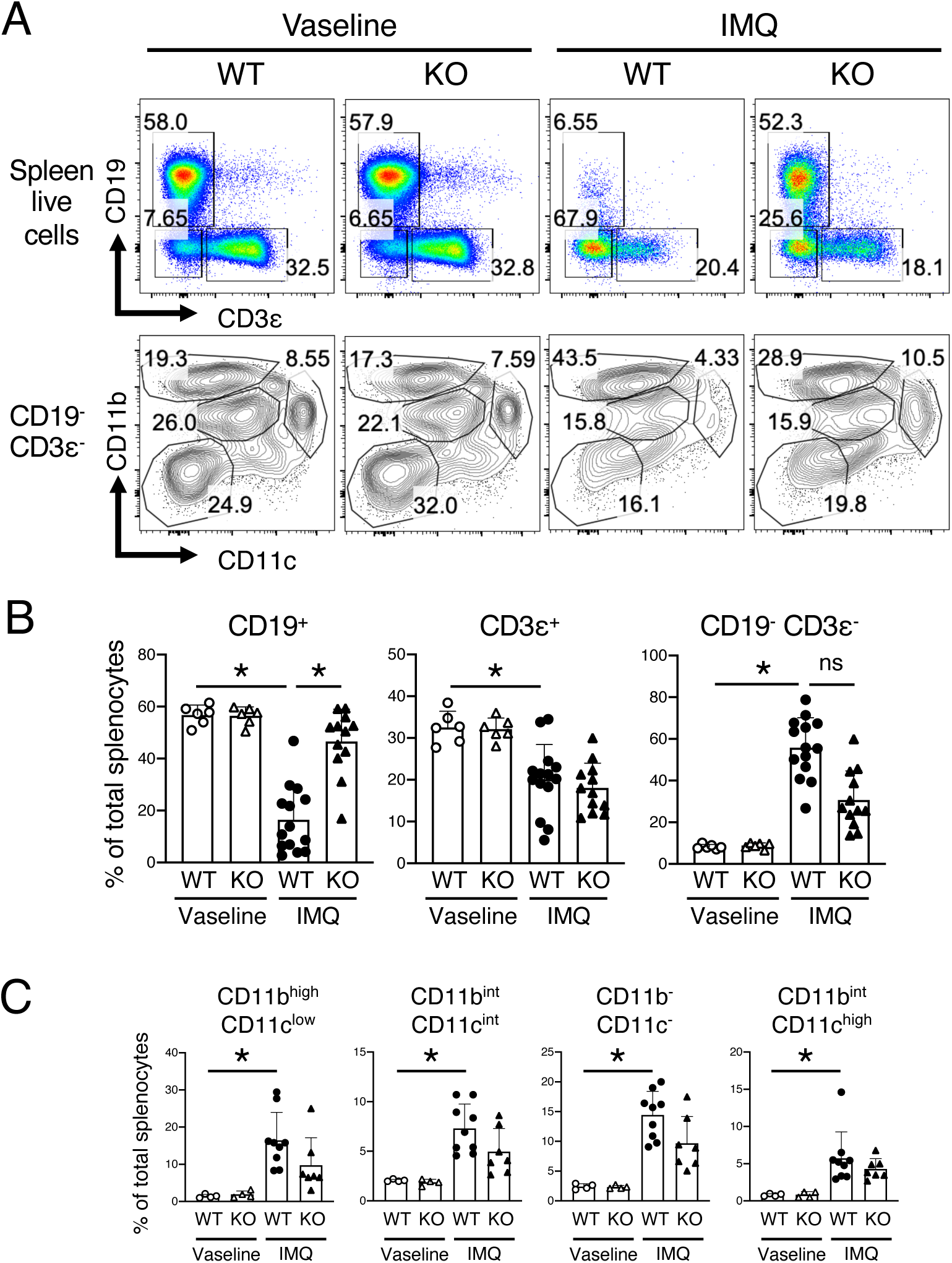
*Ebi3* deficiency partially retrieved imiquimod (IMQ)-induced changes in immune cell subsets in the spleen. IMQ or Vaseline was applied to the ears of WT or *Ebi3* knock-out (KO) mice three times a week for 8 weeks. Immune cell subsets in the spleen were analyzed by flow cytometry. (A) Representative results of CD19 and CD3ε expression (upper) and CD11b and CD11c expression in CD19^-^ CD3ε^-^ population (lower) in the splenocytes. Numbers in the plots indicate the percentage of cells in each gate. (B) Percentages of CD19^+^, CD3ε^+^, and CD19^-^ CD3ε^-^ cells in the live splenocytes from experiments shown in (A). (C) Percentage of CD11b^high^ CD11c^low^, CD11b^int^ CD11c^int^, CD11b^-^ CD11c^-^, and CD11b^int^ CD11c^high^ cells in the live splenocytes from experiments shown in (A). All data are expressed as the mean + standard deviation. \**P* < 0.05; ns: not significant; by the Kruskal-Wallis test with the Dunn’s multiple comparisons test.

The analysis of T cell subsets revealed that the percentage of CD8^+^ T cells was significantly reduced in the IMQ-treated WT mice (Supplementary Fig. 4A and B). Among CD4^+^ T cells, the naive T cells were significantly reduced, while effector/memory T cells were significantly increased after IMQ application (Supplementary Fig. 4A and C). EBI3 deficiency partially recovered the IMQ-induced disturbance in immune cell subsets (Supplementary Fig. 4A and C).

### IMQ induced extramedullary hematopoiesis in the spleen

We performed a histological analysis of the spleen to examine the histopathological changes of the enlarged spleen. We found that large numbers of giant cells were observed in the spleen of the IMQ-treated WT mice (Fig. 3A and B). Immunohistochemistry revealed that the giant cells were Factor VIII-positive, representing that the giant cells are megakaryocytes (Supplemental Fig. 5). Although the megakaryocytes were also observed in the IMQ-applied KO mice, the number of the cells was significantly decreased compared to that in IMQ-treated WT mice (Fig. 3A and B). Next, we assessed erythroblast formation in the spleen by analyzing the expression of the erythroblast marker TER-119 through flow cytometry. We found that the percentages of both TER-119^low^ (proerythroblasts) and TER-119^high^ (erythroblasts) populations were increased in the IMQ-applied WT mice and that the increase was diminished in the IMQ-treated KO mice (Fig. 3C and D).

**Figure 3.**
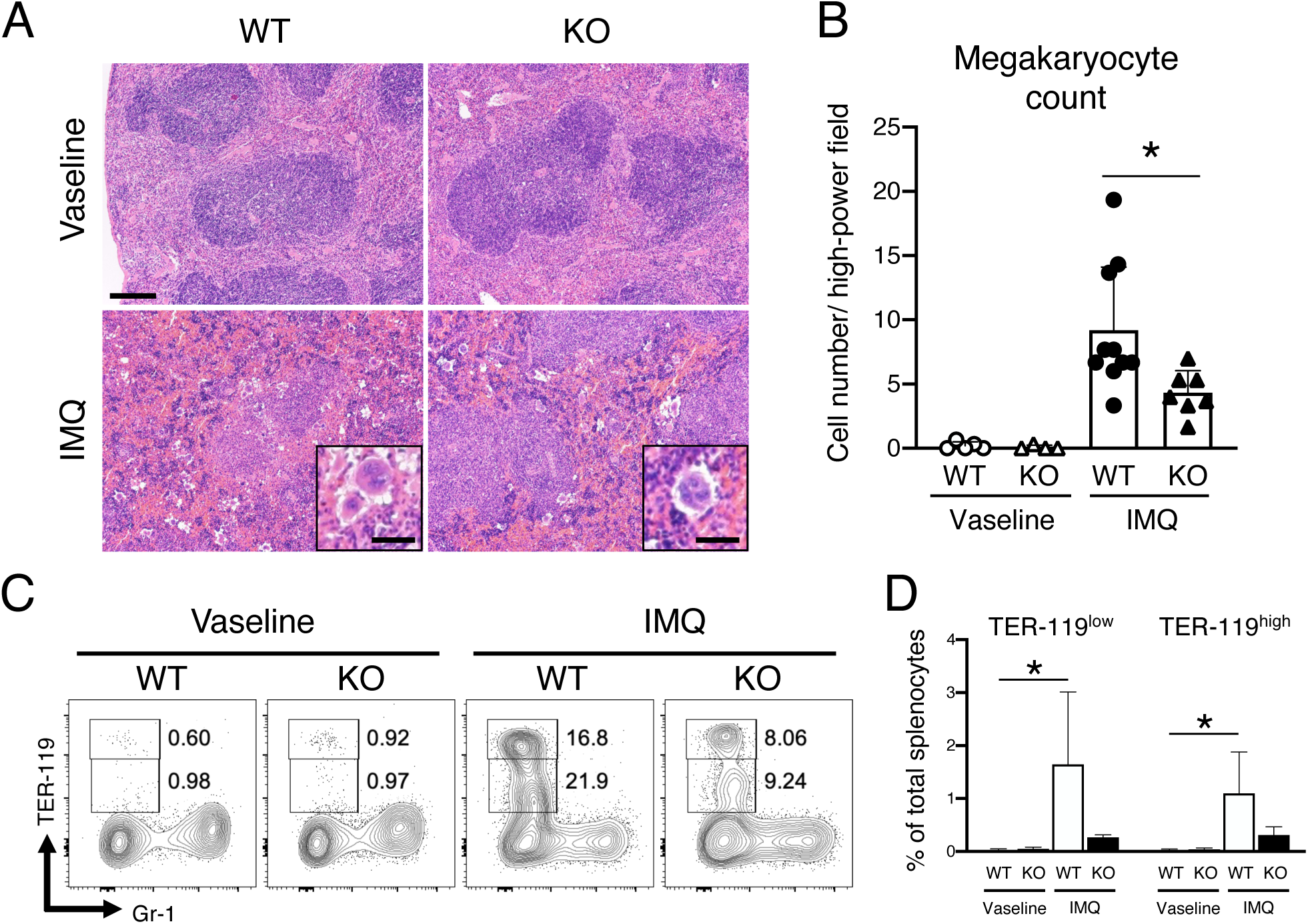
Imiquimod (IMQ)-induced extramedullary hematopoiesis was EBI3-dependent. IMQ or Vaseline was applied to the ears of wild-type (WT) or *Ebi3* knock-out (KO) mice three times a week for 8 weeks. (A) Representative images of hematoxylin and eosin staining of the spleens. The inset shows megakaryocytes. Scale bars: 200 μm for low-power fields, 40 μm for the insets. (B) Megakaryocyte counts per high-power field. (C) Representative results of Ter-119 and Gr-1 expression by flow cytometry in the splenic CD19^-^ CD4^-^ CD8^-^ CD11b^-^ cells of WT or KO mice. Numbers in the plots indicate the percentage of cells in each gate. (D) Percentages of Ter-119^low^ and Ter-119^high^ cell populations in the live splenocytes from experiments shown in (C). All data are expressed as the mean + standard deviation. \**P* < 0.05; by the Mann-Whitney test (B) or the Kruskal-Wallis test with the Dunn’s multiple comparisons test (D).

In macroscopic observation, the bone marrow in the tibia of IMQ-applied WT mice showed a whiter appearance than that of Vaseline-applied mice. The finding was not prominent in the IMQ-treated KO mice (Supplementary Fig. 6A). Tissue sections of the bone marrow of the tibia showed that cellularity was intact in the IMQ-treated WT mice compared to Vaseline-treated WT mice, except the decrease in red blood cells in the IMQ-treated WT mice (Supplementary Fig. 6B). In the IMQ-treated KO mice, cellularity was intact, and the reduced number of red blood cells was not observed (Supplementary Fig. 6B). These data indicate that IMQ-induced severe bicytopenia (anemia and thrombocytopenia) was not induced by bone marrow failure, but presumably via increased splenic function reflecting marked splenomegaly.

### IMQ treatment induced the activation in type I IFN-related genes

To examine inflammatory phenotypes of the mice after IMQ treatment, we performed multiplex analysis of cytokines in the sera of IMQ-applied WT and KO mice. We found no significant changes in the inflammatory cytokines tested, including IL-6, TNF-α, IL-1β, MCP-1, IL-2, IFN-γ, IL-4, IL-10, IL12p70, and IL-17 (Supplementary Fig. 7A). In consistent with the findings, no upregulation of *Il6*, *Tnf,* and *Il1b* genes was observed in the spleen and peripheral blood of IMQ-applied mice (Supplementary Fig. 7B and C).

To ascertain the impact of EBI3 deficiency in the gene expression, RNA-seq was performed using the RNA samples isolated from the spleen and peripheral blood leukocytes after 8 weeks IMQ application. Pooled samples from multiple mice of the same group were prepared and applied to the RNA-seq analysis. Based on the results of differential gene expression analysis and gene ontology (GO) analysis, we focused on genes whose expression varied between IMQ-applied WT and KO mice. The GO analysis revealed that genes related to the response to viral infection and type I IFN are among the top genes that are decreased in the KO peripheral blood compared to the IMQ-applied WT mice (Fig. 4A). The relative expression changes in the spleen and peripheral blood were examined among the four groups. The results demonstrated that individual IFN-related genes were upregulated in the IMQ-applied WT mice and that the increase of the genes tended to be milder in the IMQ-applied KO mice (Fig. 4B and C). We then performed a qPCR analysis using individual RNA samples to further validate the RNA-seq data that were obtained from pooled RNA samples. In the spleen, we observed an increase in the gene expression in the IMQ-applied WT mice and a relatively milder increase in the IMQ-applied KO mice, which was consistent with the RNA-seq data. In particular, the expression of *Isg15*, *Irf7*, and *Ifit1* was significantly elevated in WT mice following IMQ application (Fig. 4D). On the other hand, in peripheral blood, both WT and KO mice showed similar levels of increased gene expression after IMQ treatment (Fig. 4E).

**Figure 4.**
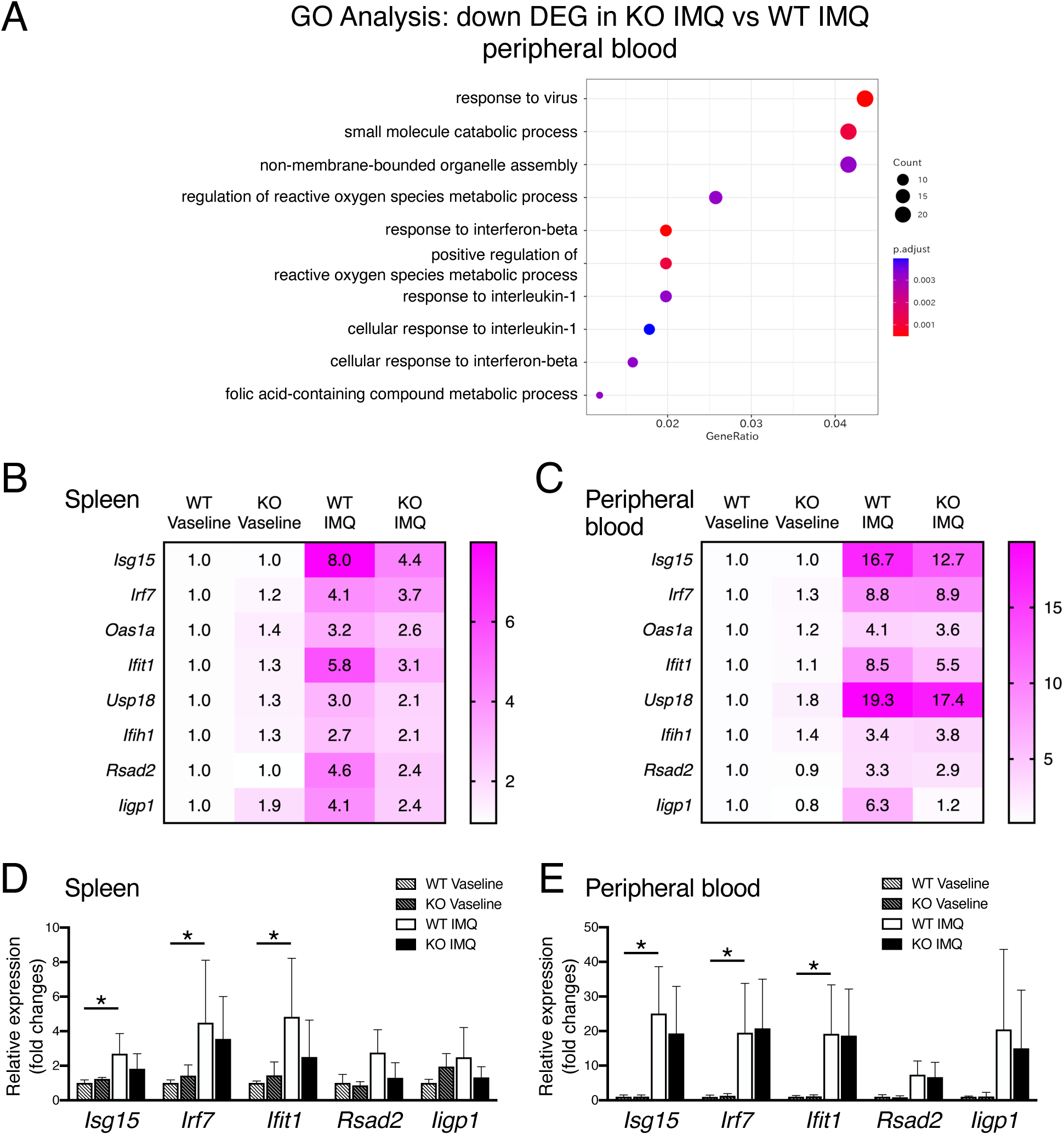
Type I interferon (IFN)-related gene activation induced by imiquimod (IMQ) is reduced by *Ebi3* knock-out (KO). (A) Gene ontology (GO) over-representation test of downregulated genes in the peripheral blood leukocytes of IMQ-treated *Ebi3* KO mice compared to those of the wild-type (WT) mice. (B, C) Heatmap showing relative TPM (transcripts per million) levels of IFN-signature genes from RNA-sequencing (RNA-seq) analysis of the spleen (B) and peripheral blood leukocytes (C). (D, E) Quantitative polymerase chain reaction analysis of genes that showed differences in RNA-seq analysis in the spleen (D) and peripheral blood leukocytes (E). Values in Vaseline-treated WT mice is set to 1. Expression was normalized to that of *Hprt* (D) or *Gapdh* (E). All data are expressed as the mean + standard deviation. \**P* < 0.05; by the Kruskal-Wallis test with the Dunn’s multiple comparisons test.

### *Ebi3*-deficiency does not affect pDC differentiation and TLR7-stimulated IFN production

Type I IFN is abundantly produced from pDCs under TLR7 stimulation (42). Since IMQ-induced type I IFN-related gene expression was decreased in the KO mice compared to WT mice, we first suspected that pDC differentiation is suppressed in the KO mice. We thus examined the proportion of pDC and cDC in the spleens of untreated WT and KO mice. We found no significant difference in the percentages of pDCs and cDCs (Fig. 5A and B). We then considered the possibility that the production of type I IFNs from pDCs is suppressed in the KO mice. To test this, we examined *Ifna* gene expression and serum IFN-α levels. *Ifna* gene expression was induced after IMQ treatment in the skin, not in the spleen. In addition, IFN-α concentration in the sera was increased after IMQ treatment (Fig. 5D). In terms of the phenotypical differences between WT and KO mice, the increased expression was comparable between WT and KO mice (Fig. 5C).

**Figure 5.**
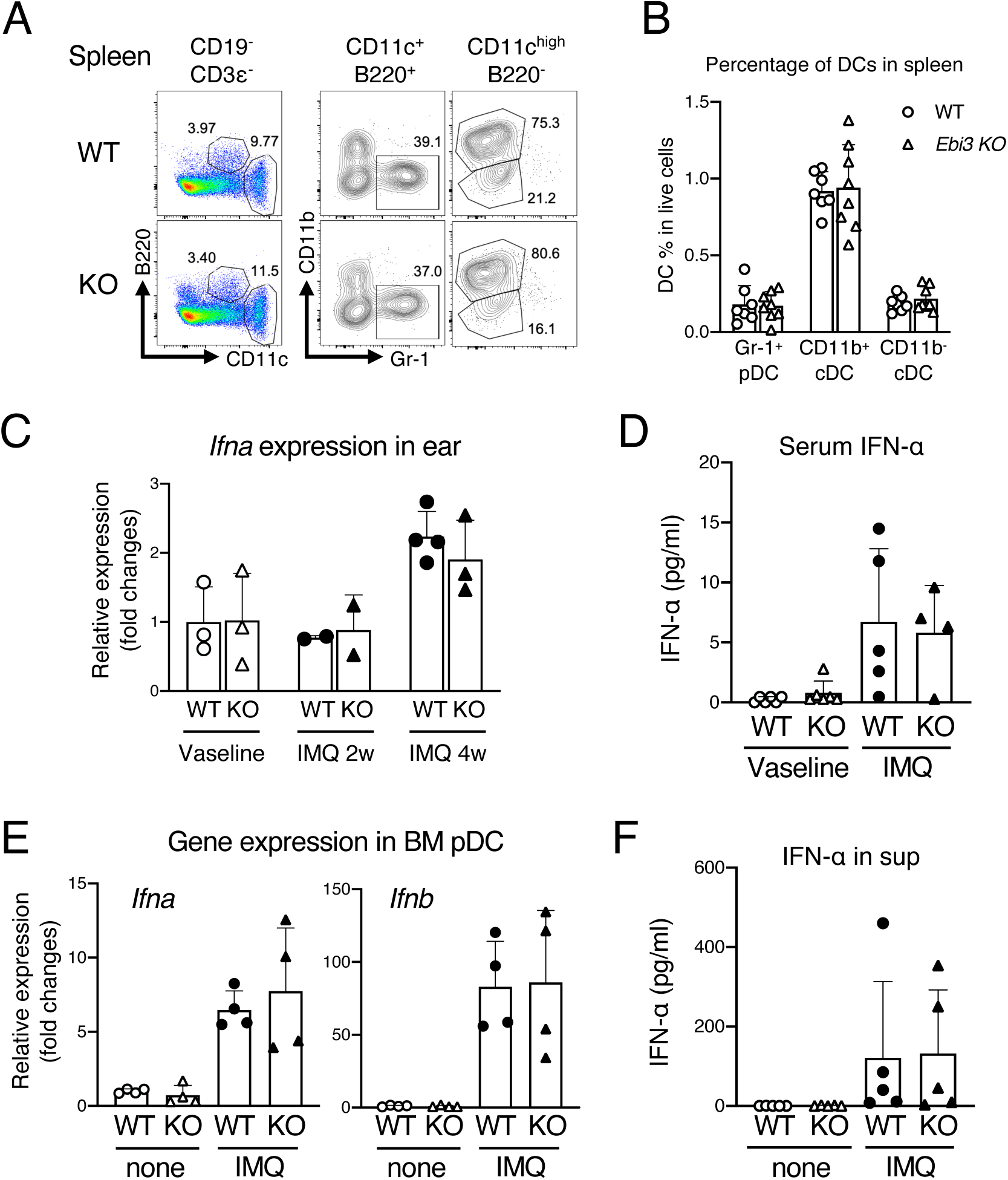
*Ebi3* deficiency did not alter the development and function of plasmacytoid dendritic cells (pDCs). (A) Representative flow cytometric analysis of the percentage of pDCs (CD11c^+^ B220^+^) and conventional dendritic cells (cDCs) (CD11c^+^ B220^-^) in the spleen of wild-type (WT) or KO mice. Numbers in the plots indicate the percentage of cells in each gate. (B) Percentages of pDCs (CD11c^+^ B220^+^ Gr-1^+^) or cDCs (CD11c^+^ B220^-^ CD11b^+^ or CD11c^+^ B220^-^ CD11b^-^) in the spleen of WT or KO mice. (C) *Infa* gene expression in ear skin sample of Vaseline- or imiquimod (IMQ)-treated WT or KO mice. Gene expression was analyzed with quantitative polymerase chain reaction (qPCR). (D) IFN-α concentration in serum of Vaseline- or IMQ-treated WT or KO mice. IFN-α concentrations were quantified using enzyme-linked immunosorbent assay (ELISA). (E, F) Bone marrow-derived pDCs were stimulated with 200 ng/ml of IMQ for 6 hours. (E) *Ifna* gene expression in the bone-marrow derived pDCs. Expression levels were analyzed by qPCR and normalized to that of *Hprt.* (F) The concentrations of IFN-α in the culture supernatants. All data are expressed as the mean + standard deviation.

For further analysis, we assessed pDCs differentiation and activation in the bone marrow-derived pDCs culture from WT and KO mice. Bone marrow cells were cultured in the presence of hFLT3L for one week and analyzed by flow cytometry. The results indicated that the percentages of CD11c^+^ B220^+^ pDCs were comparable (WT, 20.1–25.1%; KO, 18.6–25.4%) (Supplementary Fig. 8A and B). The expression levels of cell surface MHC class II were also comparable between WT and KO DCs (Supplementary Fig. 8C). The induction of the type I IFN genes was also similar between WT and KO DCs (Fig. 5E). These findings suggest that pDCs and type I IFN are less likely to attribute to the blunted induction in type I IFN-related genes of the IMQ-treated KO mice.

### IL-27 was induced by IMQ and was likely to attribute to type I IFN-related gene induction

Although increased expression of type I IFN-related genes was observed in IMQ-applied WT and decreased in KO, the type I IFN production was not altered between WT and KO mice. These findings suggest that factors other than type I IFN induce type I IFN-related genes in the spleen in correspondence with EBI3 expression. Regarding the possible factor, recent studies have shown that IL-27 induces the expression of type I IFN-related genes independently of type I IFN (13). In our study, we hypothesized that the differences in the presence or absence of IL-27 caused the phenotypic differences between WT and KO mice, including the blunted expression levels of type I IFN genes in the KO mice. To this end, we initially examined whether IMQ application induced the expression of the *Il27* and *Ebi3* genes in WT mice. We found that IMQ application induced both *Il27* and *Ebi3* expression in the spleen and peripheral blood in WT mice (Fig. 6A).

**Figure 6.**
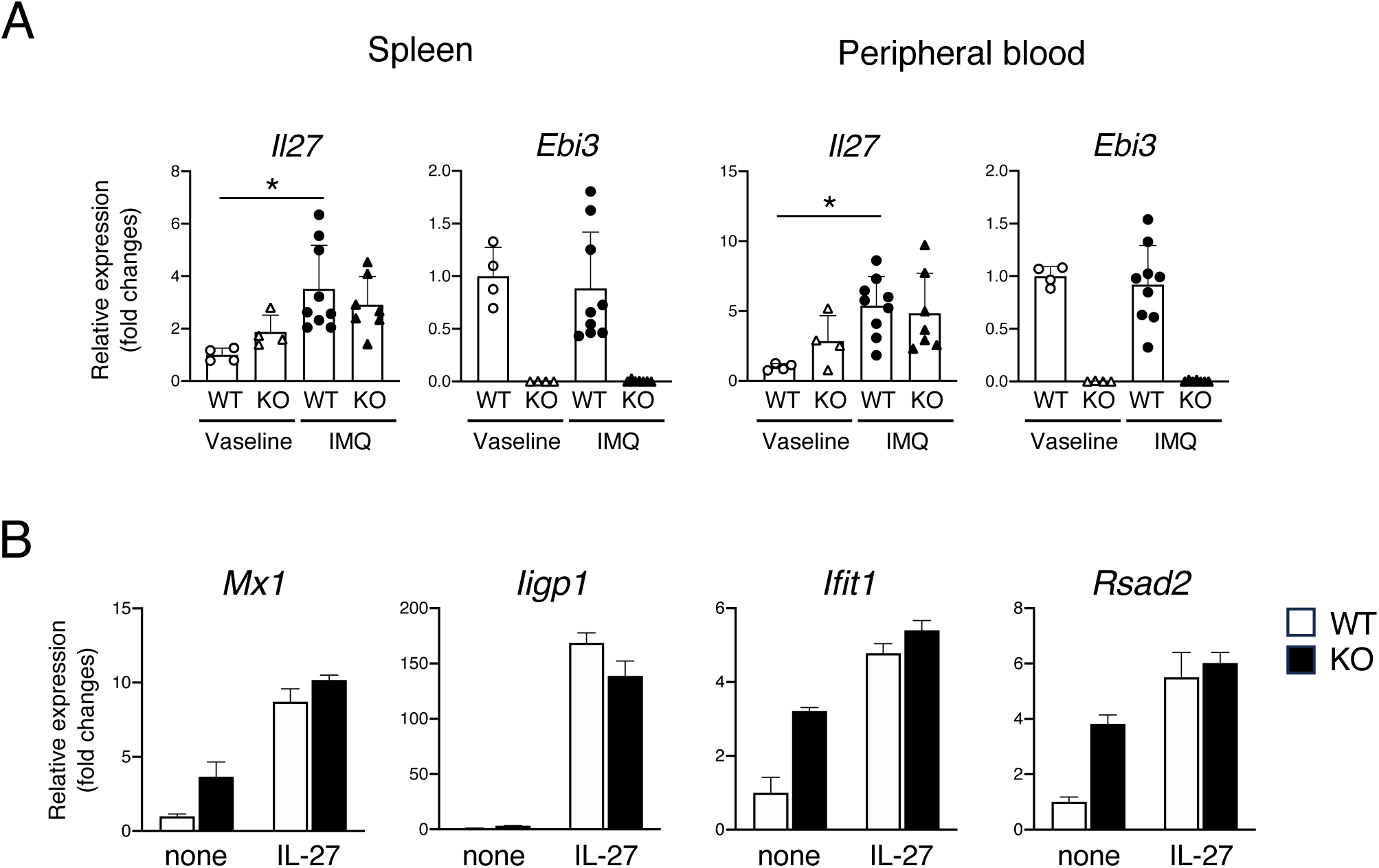
IL-27 induced by IMQ application upregulates type I IFN-related genes. (A) *Il27* and *Ebi3* gene expression in spleen and peripheral blood cells of Vaseline- or IMQ-treated WT or KO mice. (B) Expression analysis of type I IFN-related genes in bone marrow-derived macrophages stimulated with 100 ng/ml of IL-27. All data are expressed as the mean + standard deviation. \**P* < 0.05; by the Kruskal-Wallis test with the Dunn’s multiple comparisons test.

We next investigated whether recombinant IL-27 can induce type I IFN-related genes in the bone marrow-derived macrophage culture. The results demonstrated that IL-27 treatment induced type I IFN-related genes, such as *Mx1*, *Iigp1*, *Ifit1* and *Rsad2*, in the WT and KO macrophages (Fig. 6B). These findings indicate that IL-27 is produced the WT mice in response to IMQ and that IL-27 has capacity to induce type I IFN-related genes as reported (Ref). Considering the findings, EBI3 deficiency, which means the absence of IL-27, results in a reduction in the type I IFN-related gene expression for the IL-27-dependent portion.

## Discussion

In this study, we explored the role of EBI3 in the TLR7-mediated splenomegaly and cytopenia. We demonstrated the following findings. 1) Repeated TLR7 stimulation induces splenomegaly and severe bicytopenia (anemia and thrombocytopenia), associated with extramedullary hematopoiesis in the spleen, 2) Repeated TLR7 stimulation increases myeloid cells in the enlarged spleen. 3) Repeated TLR7 stimulation upregulates Type I IFN-related gene expression in the spleen and peripheral blood, 4) *Ebi3* deficiency partially retrieved the IMQ-induced above findings, 5) *Ebi3* deficiency does not alter the differentiation and functions of pDCs, 6) IL-27, a heterodimer of EBI3 and IL-27p28, induces type I IFN-related genes. Based on the findings, we hypothesize that impaired IL-27 production by EBI3 deficiency leads to suppressed type I IFN-related gene expression and reduced splenomegaly under repeated TLR7 stimulation.

Marked splenomegaly was observed in the IMQ-treated mice, and myeloid cells were proliferated in the enlarged spleen, as shown in Fig. 2. *In vitro* culture experiments and analysis of overexpressing mice have shown that IL-27 also acts on HSCs and promotes their proliferation and differentiation into myeloid cells (11, 12). The IMQ-treated mice induced the production of both type I IFN and IL-27, suggesting that proliferation and differentiation into myeloid cells progressed in response to IMQ. In contrast, *Ebi3* KO mice do not produce IL-27, indicating that only type I IFN affects HSCs. The weakening of the differentiation-promoting effect on myeloid cells in KO mice may contribute to the reduction of splenomegaly in the IMQ-treated KO mice.

One of the possible mechanisms is that IL-27 production in the spleen and peripheral blood cells by IMQ application induces type I IFN-related genes and activates various immune cells, including myeloid cells, T cells, and HSCs. It has recently been reported that IL-27 stimulates macrophages and induces type I IFN-related genes (43), and we also confirmed in this study that IL-27 stimulation induces type I IFN-related genes in bone marrow-derived macrophages (Fig. 6C). Therefore, IL-27 is likely to induce splenomegaly through stimulation of myeloid cells and induction of type I IFN-related gene expression.

IL-27 has been reported to activate T cells and induce Th1 differentiation (8–10). We have found that IMQ application activates splenic CD4^+^ cells in WT mice, which was blunted in the KO mice (Supplementary Fig. 4). Since IL-27 stimulates T cells to induce expression of type I IFN-related genes (44), it is possible that the T cells are among the cells expressing type I IFN-related genes in response to IL-27.

Although *Ebi3* deficiency retrieved the IMQ-mediated splenomegaly, bicytopenia, and IFN-related gene expression, the suppressive effect of *Ebi3* deficiency was not complete. These findings suggest that the IMQ-induced pathological changes are partly mediated by some factors other than cytokines constituted by EBI3 and EBI3-related pathways. IFN-λ and its related pathways might be involved in the IMQ-induced pathological changes independently of EBI3. Goel et al. have shown that splenomegaly, leukocytosis, anemia, and thrombocytopenia are relieved by IFN-λ receptor deficiency (29). IFN-λ has also been shown to induce type I IFN-related gene expression. Whether the IFN-λ pathway and the EBI3-mediated pathways are independent or potentially related needs further investigation.

EB virus infection, named infectious mononucleosis, has been known to cause splenomegaly. Splenic rupture associated with marked splenomegaly is recognized as a critical complication of infectious mononucleosis (18). It is speculated that continuous TLR7 stimulation by the viral infection leads to the production of type I IFN and IL-27, which in turn contributes to the development of splenomegaly. Inhibition of the cytokines might be a therapeutic option for high-risk patients for splenic rupture by reducing enlargement of the spleen.

Based on our findings in this study, we consider that EBI3 is involved in the pathogenesis of splenomegaly and bicytopenia via IL-27. Since EBI3 has multiple functions in immune regulation, we might need to consider other functions of EBI3. EBI3 binds to IL-12p35 and IL-23p19 and functions as heterodimer cytokines named IL-35 and IL-39, respectively (5). EBI3 also serves as a molecular chaperone in conjunction with calnexin in CD4^+^ T cells during inflammation (45). Additionally, EBI3 modulates soluble IL-6 receptor-like function (46) and functions as an inhibitory cytokine with IL-12p40 (47). The EBI3-mediated immune regulation might be involved in the IMQ-induced pathological changes observed in this study. Further analysis is needed to obtain a complete picture of the action of EBI3.

In conclusion, EBI3, as a component of IL-27, regulates splenomegaly and bicytopenia that occur upon sustained TLR7 stimulation. Further elucidation of the roles of EBI3 and IL27 in immune cells and hematopoietic cells will lead to the development of novel therapies for splenomegaly and severe cytopenia associated with chronic viral infection.

## List of abbreviations

EBI3: Epstein-Barr virus induced 3
IMQ: imiquimod
IFN: niterferon
HSC: hematopoietic stem cell
TLR: toll-like receptor
pDC: plasmacytoid dendritic cell
TNF: tumor-necrosis factor
IL: interleukin
Th: helper T cell
WT: wild type
KO: knock-out
CRISPR: clustered regularly interspaced short palindromic repeats
Cas9: CRISPR associated protein 9
qPCR: quantitative polymerase chain reaction
WBC: white blood cell
RBC: red blood cell
PLT: platelet
7AAD: 7-amino-actinomycin D
hFLT3L: human FMS-like tyrosine kinase 3 ligand
ELISA: enzyme-linked immunosorbent assay
PVDF: polyvinylidene difluoride
RNA-seq: RNA-sequencing
GO: gene ontology

## Acknowledgments

We thank Keiko Watanabe and Noriko Miyake (Department of Immunology and Molecular Genetics, Kawasaki Medical School) for their technical assistance, Drs. Tatsushi Shiomi and Hirotake Nishimura (Department of Pathology, Kawasaki Medical School) for their valuable comments on histological analysis, and Drs. Tatsuo Ito and Yurika Shimizu (Department of Hygiene, Kawasaki Medical School) for their support of the multiplex analysis. We are also indebted to the staff of the Central Research Institute of Kawasaki Medical School.

## Authors’ contributions

M.Is. performed most experiments, interpreted results, and drafted and edited the manuscript. Y.S. performed experiments, interpreted the data, and reviewed the manuscript. D.T., Y.M., and S.M. interpreted the data and reviewed the manuscript. M.In. generated *Ebi3* KO mice and reviewed the manuscript. T.M. directed the project and edited the manuscript. All authors read and approved the final version of the manuscript.

## Availability of data and materials

All data generated or analyzed during this study are included in this published article and its supplementary information files.

## Declarations

## Ethics approval and consent to participate

All animal experiments were approved by the Institutional Safety Committee for Recombinant DNA Experiments (20-49, 0004-00) and the Institutional Animal Care and Use Committee of Kawasaki Medical School (22-072, 23-092).

## Consent for publication

Not applicable

