## Supplementary Table for "Epstein-Barr virus induced 3 attributes to TLR7-mediated splenomegaly and bicytopenia"

**Supplemental Table 1. qPCR primers used in this study**

| Gene | Sequence | | Amplicon  size (bp) |
| --- | --- | --- | --- |
| *Ebi3* | Forward:  Reverse: | GCTCCCCTGGTTACACTGAA  ACGGGATACCGAGAAGCAT | 79 |
| *Hprt* | Forward:  Reverse: | TCCTCCTCAGACCGCTTTT  CCTGGTTCATCATCGCTAATC | 90 |
| *Gapdh* | Forward:  Reverse: | ATCAAGAAGGTGGTGAAGCA  GACAACCTGGTCCTCAGTGT | 75 |
| *Isg15* | Forward:  Reverse: | AGTCGACCCAGTCTCTGACTCT  CCCCAGCATCTTCACCTTTA | 143 |
| *Irf7* | Forward:  Reverse: | CTTCAGCACTTTCTTCCGAGA  TGTAGTGTGGTGACCCTTGC | 68 |
| *Ifit1* | Forward:  Reverse: | TACAGCAACCATGGGAGAGAA  CAAGGAACTGGACCTGCTCTG | 149 |
| *Rsad2* | Forward:  Reverse: | AGCAATGGCAGCCTTATCCG  TTCCTTGACCACGGCCAATC | 124 |
| *Iigp1* | Forward:  Reverse: | CGATAGTAGTGTGCTCAATGTTGC  CCATGGTTACCTCCACCACC | 134 |
| *Ifna* | Forward:  Reverse: | CCTGAGA(A/G)AGAAGAAACACAGCC  GGCTCTCCAGA(C/T)TTCTGCTCTG | 64 |
| *Ifnb* | Forward:  Reverse: | CTGGCTTCCATCATGAACAA  AGAGGGCTGTGGTGGAGAA | 73 |
| *Il27* | Forward:  Reverse: | TGTCCACAGCTTTGCTGAATC  AAGTGTGGTAGCGAGGAAGC | 148 |
| *Mx1* | Forward:  Reverse: | Ctggataccaagtaaacatcctga  tgcctgcacagattattcaca | 75 |
