## Supplementary Figures for "Epstein-Barr virus induced 3 attributes to TLR7-mediated splenomegaly and bicytopenia"

### Supplementary Figure 1

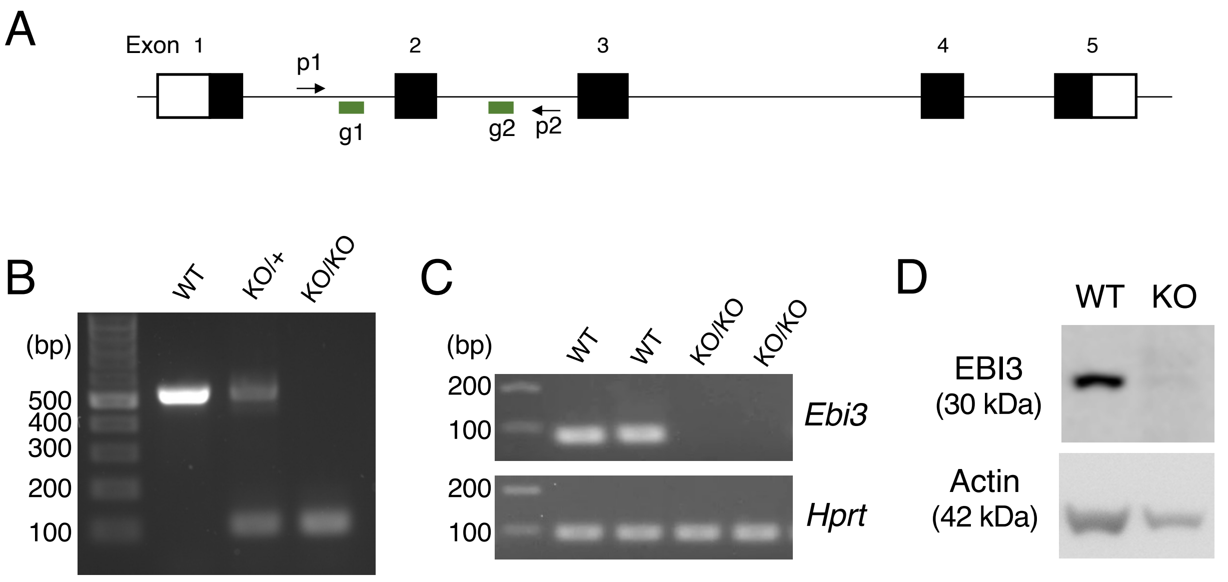


**Supplementary Figure 1.** Generation of *Ebi3*-deficient (KO) mice by CRISPR/Cas9 system. (A) Schematic structure of *Ebi3* gene. Black boxes indicate coding regions. g1 and g2 indicate guide RNAs. p1 and p2 indicate primers for genotyping. The exon 2 was deleted after genome editing. (B) Polymerase chain reaction (PCR) analysis of the genotypes of the *Ebi3*-deficient mice. (C) Reverse transcription (RT)-PCR analysis. RNA samples were isolated from the spleen of the indicated mice, and cDNA were transcribed by RT reaction. *Ebi3* mRNA expression was evaluated by RT-PCR. (D) Western blot analysis. Protein samples were collected from the spleens of the indicated mice. EBI3 protein was detected by anti-EBI3 antibody. CRISPR, clustered regularly interspaced short palindromic repeats; Cas9, CRISPR-associated protein 9; bp, base pair.

Supplementary Figure 2

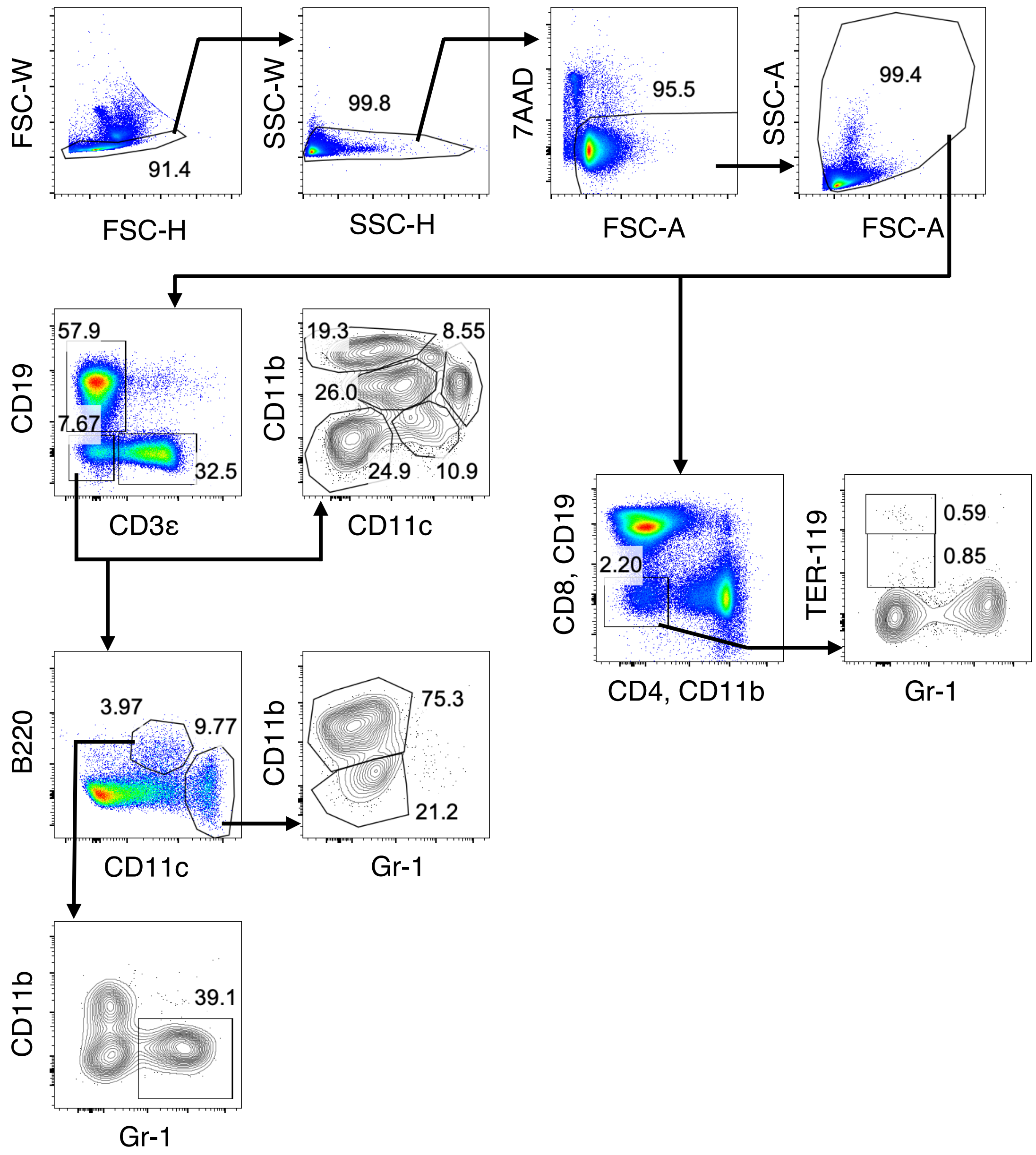


**Supplementary Figure 2.** Gating strategies for the flow cytometry analysis of the splenocytes for Fig. 2A (middle row of the left), Fig. 3C (middle row of the right), and Fig. 5A (bottom left).

Supplementary Figure 3

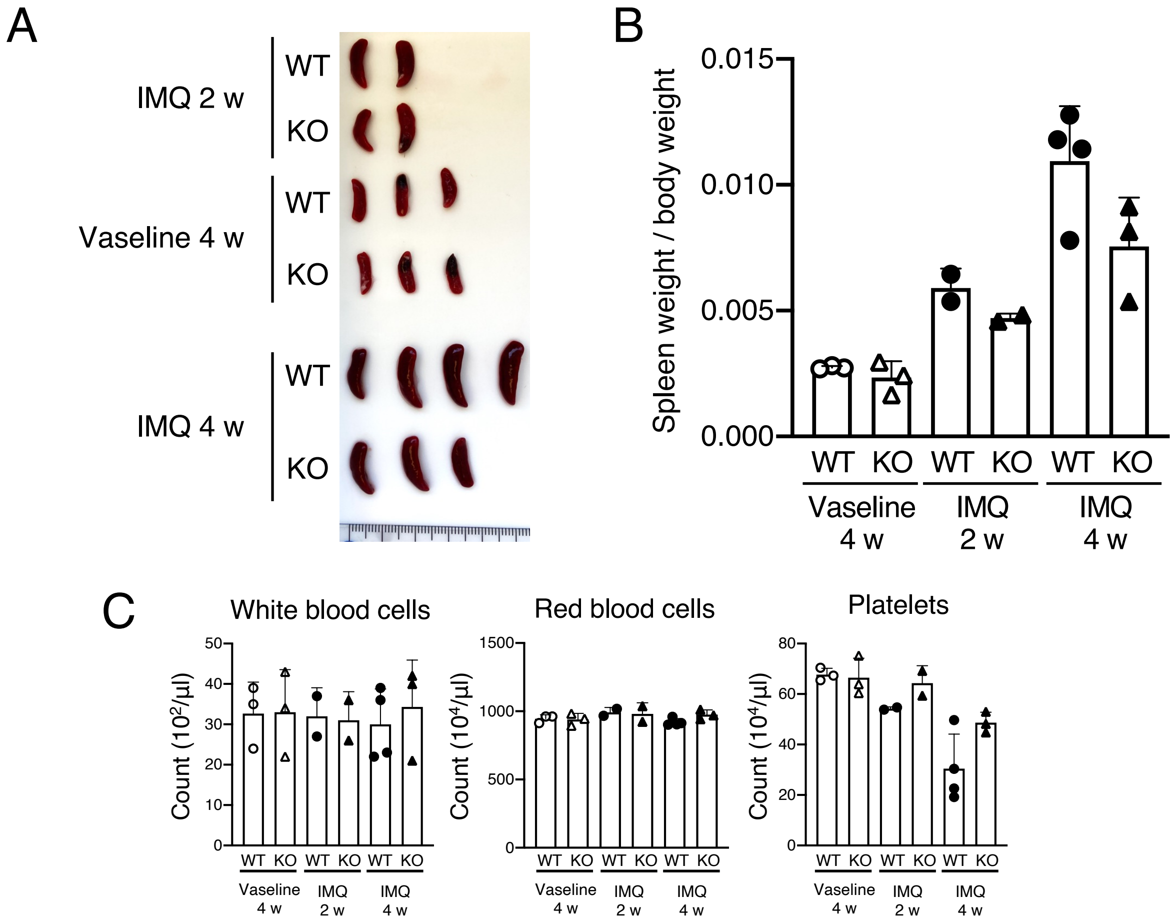


**Supplementary Figure 3.** Phenotypic analysis of the spleen and peripheral blood of imiquimod (IMQ)-treated wild-type (WT) or *Ebi3* knock-out (KO). IMQ or Vaseline was applied to the ears of WT or *Ebi3* KO mice three times a week for 2 or 4 weeks. (A) Representative macroscopic images of the spleens. (B) Spleen weight relative to body weight of WT or KO mice. (C) White blood cell count, red blood cell count, and platelet count in the peripheral blood of the mice. All data are expressed as the mean + standard deviation.

### Supplementary Figure 4

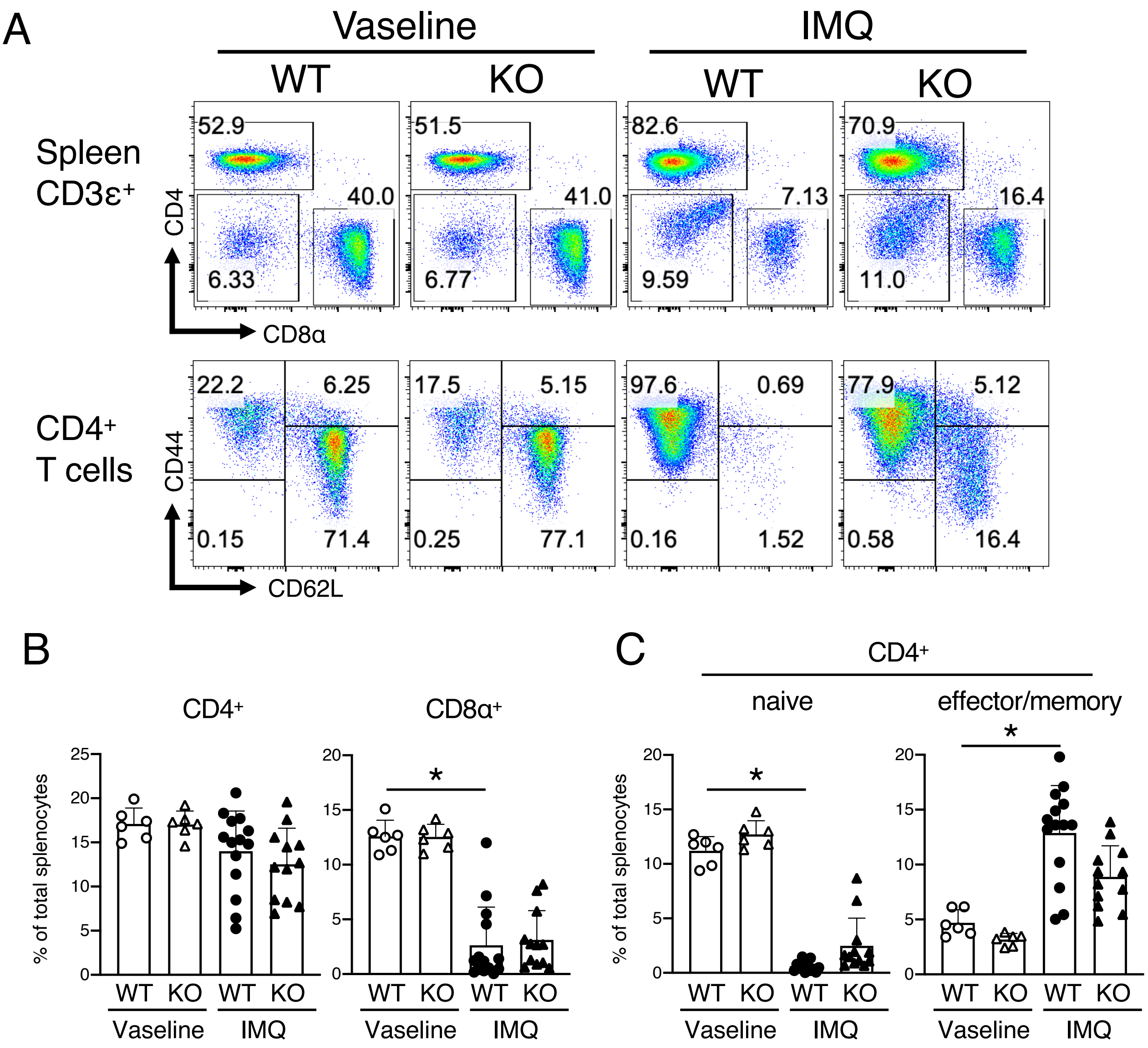


**Supplementary Figure 4.** *Ebi3* deficiency reduced imiquimod (IMQ)-induced $CD4^+$ effector/memory T cells in the spleens compared to wild-type (WT). IMQ or Vaseline was applied to the ears of WT or *Ebi3* knock-out (KO) mice three times a week for 8 weeks. (A) Representative results of flow cytometric analysis of CD4 and CD8 expression in the splenic $CD3\varepsilon^+$ cells (upper) or CD44 and CD62L expression in the splenic $CD4^+$ T cells (lower) of WT or *Ebi3* KO mice. Numbers in the plots indicate the percentage of cells in each gate. (B) Percentages of the $CD4^+$ T and $CD8^+$ T cells in the live splenocytes from experiments shown in (A). (C) Percentages of the $CD4^+$ naive T ($CD44^{low}$ $CD62L^+$) and effector/memory T ($CD44^{high}$ $CD62L^-$) cells in the spleen. All data are expressed as the mean + standard deviation. $^*P < 0.05$ by the Kruskal-Wallis test with the Dunn's multiple comparisons test.

### Supplementary Figure 5

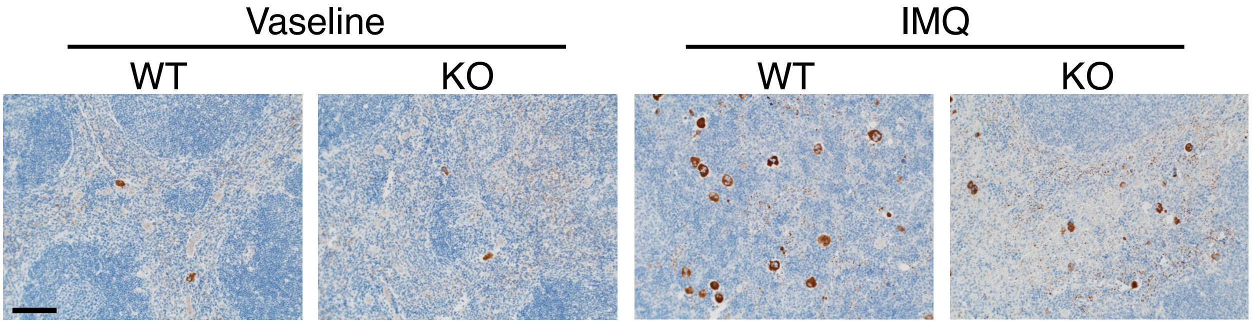


**Supplementary Figure 5.** Representative images of immunohistochemistry of spleens of wild-type (WT) or *Ebi3* knock-out (KO) mice. Sections were stained with anti-factor VIII antibody. Scale bar: 100 µm.

Supplementary Figure 6

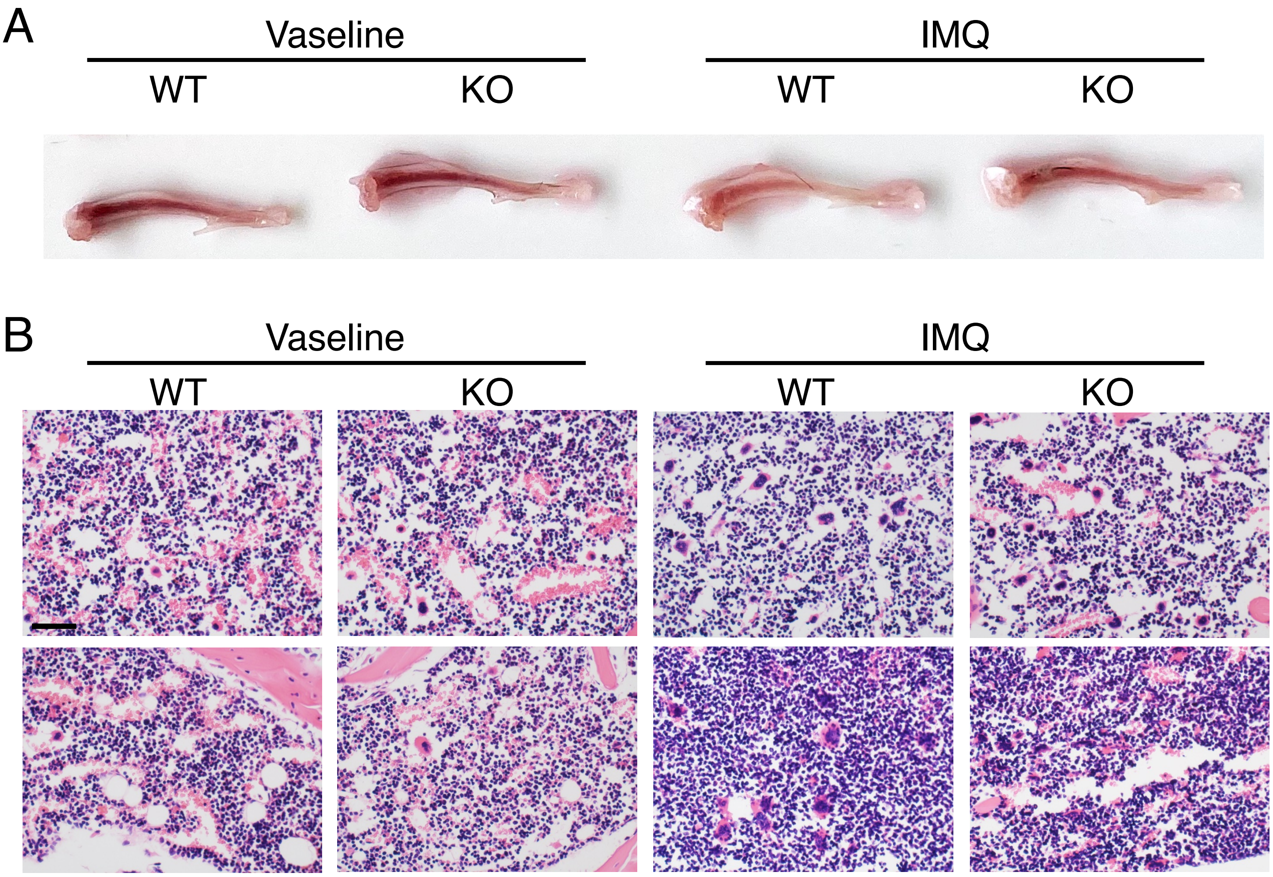


**Supplementary Figure 6.** Bone marrow phenotypes of wild-type (WT) or *Ebi3* knock-out (KO) mice after imiquimod (IMQ) application. IMQ or Vaseline was applied to the ears of WT or *Ebi3* KO mice three times a week for 8 weeks. (A) Representative photographs of tibias from WT or *Ebi3* KO mice. (B) Representative images of hematoxylin and eosin staining of bone marrows of wild-type or *Ebi3* KO mice. Scale bar: 50 µm.

### Supplementary Figure 7

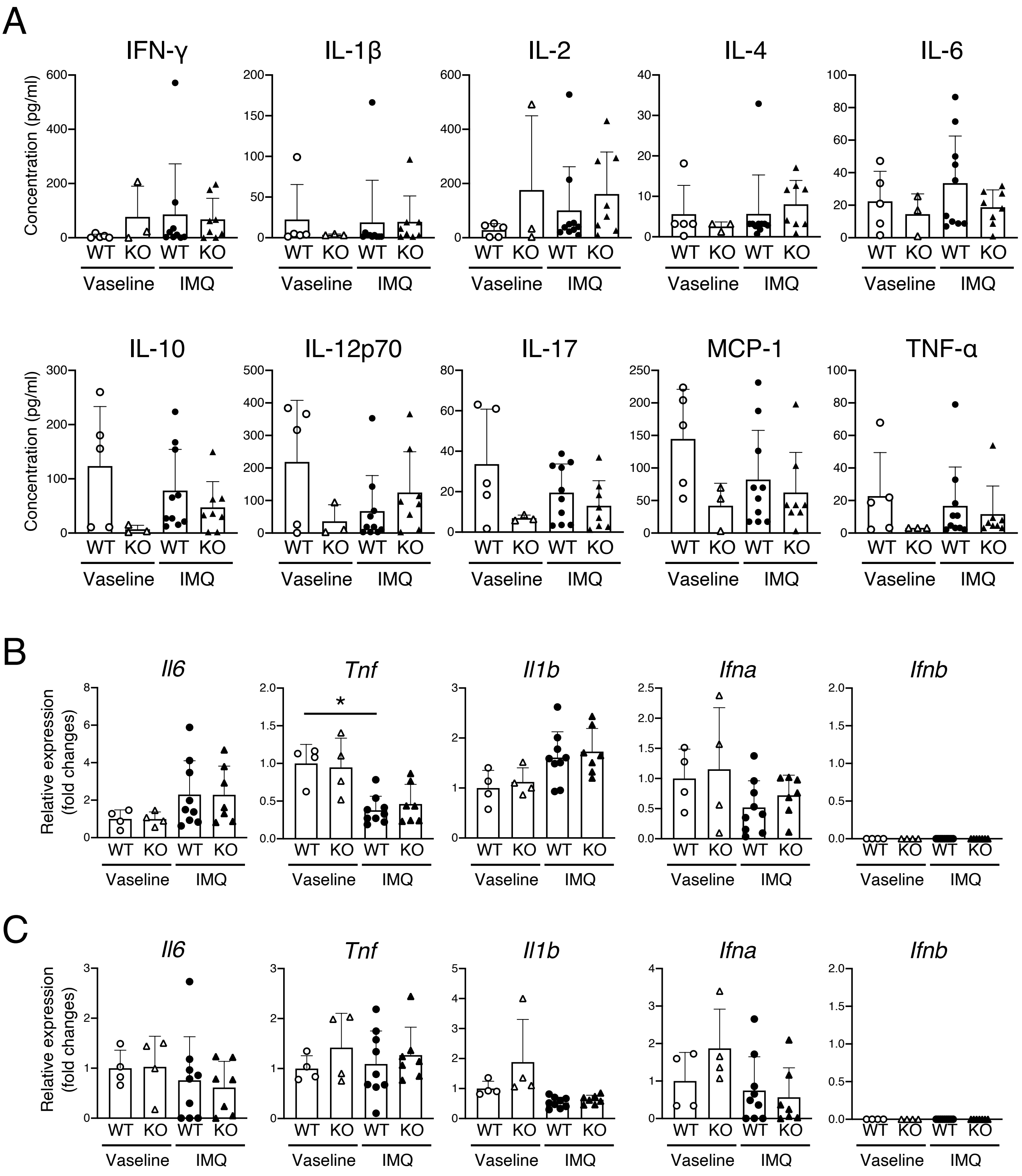


**Supplementary Figure 7.** (A) Serum cytokine levels of wild-type (WT) or *Ebi3* knock-out (KO) mice. IMQ or Vaseline was applied to the ears of WT or KO mice three times a week for 8 weeks. Serum cytokine levels were determined by multiplex cytokine assay. (B, C) Gene expression in the spleen (B) or peripheral blood (C) of the indicated mice. All data are expressed as the mean + standard deviation. *$P < 0.05$ by the Kruskal-Wallis test with the Dunn's multiple comparisons test.

### Supplementary Figure 8

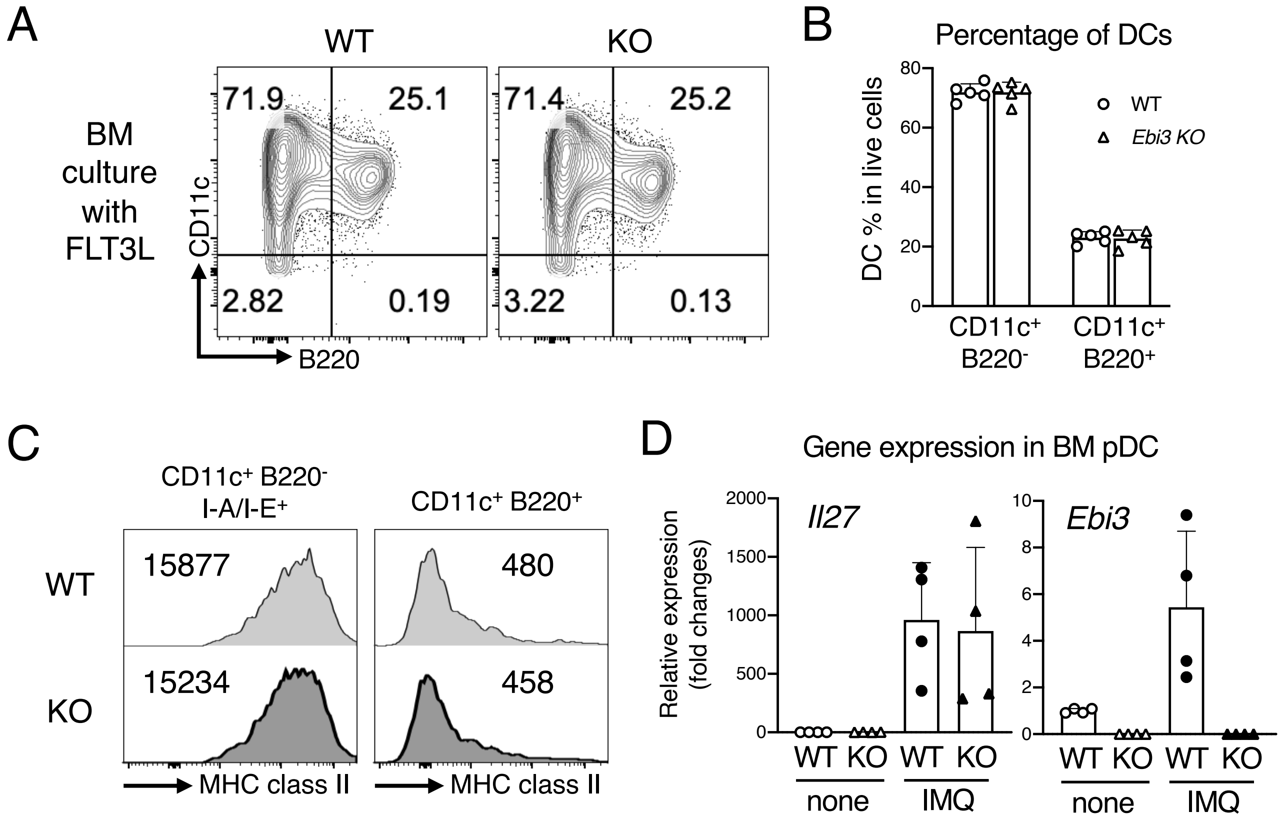


**Supplementary Figure 8.** (A-D) Bone marrow cells were collected from WT or *Ebi3* KO mice and cultured in the presence of human fms-like tyrosine kinase 3 ligand (hFLT3L) for 1 week. (A) Representative results of flow cytometric analysis of CD11c and B220 expression. Numbers in the plots indicate the percentage of cells in each quadrant gate. (B) Percentage of pDCs (CD11c$^{+}$ B220$^{+}$) and cDCs (CD11c$^{+}$ B220$^{-}$) in the cultured cells. (C) Representative plots of cell surface MHC class II expression of pDC and cDC in the cultured cells. (D) *Il27* and *Ebi3* gene expression. Bone marrow-derived pDCs were stimulated with 200 ng/ml of IMQ for 6 hours. *Il27* and *Ebi3* gene expression was analyzed by qPCR. All data are expressed as the mean + standard deviation.
